## Supplementary material for "Selective sorting of ancestral introgression in maize and teosinte along an elevational cline": S3 Table

**S3 Table. Spearman's rank correlation between genomewide admixture proportions (NGSAdmix) and recombination rate (or coding bp per cM) quintiles**

| group | feature | ancestry | Spearman's $\rho$ | 2.5% | 97.5% |
|---|---|---|---|---|---|
| sympatric mexicana | recombination rate (cM/Mb) | maize | -1.00 | -1.00 | -0.90 |
| sympatric mexicana | recombination rate (cM/Mb) | mexicana | 1.00 | 0.80 | 1.00 |
| sympatric mexicana | recombination rate (cM/Mb) | parviglumis | 0.00 | -0.90 | 0.90 |
| sympatric maize | recombination rate (cM/Mb) | maize | -1.00 | -1.00 | -0.85 |
| sympatric maize | recombination rate (cM/Mb) | mexicana | 1.00 | 0.80 | 1.00 |
| sympatric maize | recombination rate (cM/Mb) | parviglumis | 0.70 | 0.10 | 0.95 |
| reference parviglumis | recombination rate (cM/Mb) | maize | -0.50 | -1.00 | -0.35 |
| reference parviglumis | recombination rate (cM/Mb) | mexicana | 0.10 | -0.95 | 0.50 |
| reference parviglumis | recombination rate (cM/Mb) | parviglumis | 0.50 | 0.30 | 1.00 |
| reference mexicana | recombination rate (cM/Mb) | maize | -0.70 | -1.00 | -0.40 |
| reference mexicana | recombination rate (cM/Mb) | mexicana | 0.50 | -0.30 | 0.80 |
| reference mexicana | recombination rate (cM/Mb) | parviglumis | -0.10 | -0.70 | 0.50 |
| reference maize | recombination rate (cM/Mb) | maize | 0.30 | -0.25 | 0.70 |
| reference maize | recombination rate (cM/Mb) | mexicana | 0.00 | -0.70 | 1.00 |
| reference maize | recombination rate (cM/Mb) | parviglumis | -0.30 | -0.70 | 0.30 |
| sympatric mexicana | gene density (coding bp/cM) | maize | 1.00 | 0.85 | 1.00 |
| sympatric mexicana | gene density (coding bp/cM) | mexicana | -1.00 | -1.00 | -0.80 |
| sympatric mexicana | gene density (coding bp/cM) | parviglumis | -0.70 | -1.00 | 0.90 |
| sympatric maize | gene density (coding bp/cM) | maize | 1.00 | 0.90 | 1.00 |
| sympatric maize | gene density (coding bp/cM) | mexicana | -1.00 | -1.00 | -0.90 |
| sympatric maize | gene density (coding bp/cM) | parviglumis | -0.70 | -0.90 | 0.15 |
| reference parviglumis | gene density (coding bp/cM) | maize | 0.30 | 0.00 | 0.90 |
| reference parviglumis | gene density (coding bp/cM) | mexicana | 0.70 | -0.30 | 1.00 |
| reference parviglumis | gene density (coding bp/cM) | parviglumis | -0.60 | -0.90 | -0.05 |
| reference mexicana | gene density (coding bp/cM) | maize | 0.90 | 0.30 | 0.95 |
| reference mexicana | gene density (coding bp/cM) | mexicana | -0.40 | -0.90 | 0.35 |
| reference mexicana | gene density (coding bp/cM) | parviglumis | 0.30 | -0.60 | 0.90 |
| reference maize | gene density (coding bp/cM) | maize | -0.90 | -1.00 | -0.10 |
| reference maize | gene density (coding bp/cM) | mexicana | 0.10 | -0.80 | 0.80 |
| reference maize | gene density (coding bp/cM) | parviglumis | 0.90 | 0.10 | 0.90 |
