## Supplementary material for "Selective sorting of ancestral introgression in maize and teosinte along an elevational cline": S4 Table

**S4 Table. *Mexicana* ancestry by elevation and recombination rate quintile.** Best-fitting linear models for ancestry proportion predicted by an elevation by recombination rate interaction: *mexicana* ancestry $\sim$ elevation + r + elevation*r. Here, r is the recombination rate quintile, treated as numeric [0-4]. This model only uses ancestry estimates for sympatric individuals and is fit separately for maize and *mexicana* samples.

| group | term | estimate | std.error | statistic | p.value |
|---|---|---|---|---|---|
| sympatric maize | intercept | -0.147 | 0.028 | -5.307 | 1.55E-07 |
| sympatric maize | elevation (km) | 0.099 | 0.013 | 7.630 | 8.78E-14 |
| sympatric maize | r quintile | -0.114 | 0.011 | -10.034 | 4.54E-22 |
| sympatric maize | elevation*r quintile | 0.077 | 0.005 | 14.484 | 3.27E-41 |
| sympatric mexicana | intercept | 0.045 | 0.034 | 1.325 | 1.86E-01 |
| sympatric mexicana | elevation (km) | 0.366 | 0.016 | 23.252 | 7.62E-94 |
| sympatric mexicana | r quintile | 0.086 | 0.014 | 6.242 | 6.67E-10 |
| sympatric mexicana | elevation*r quintile | -0.031 | 0.006 | -4.832 | 1.59E-06 |
