## Supplementary material for "Selective sorting of ancestral introgression in maize and teosinte along an elevational cline": S5 Table

**S5 Table. Spearman's rank correlation between mean local ancestry and recombination rate (or coding bp per cM) at 1 cM genomic window resolution.** Confidence intervals are constructed using the percentile method and 10,000 bootstrap replicates created by randomly re-sampling 1 cM windows within quintiles.

| group | ancestry | feature | Spearman's $\rho$ | 2.5% | 97.5% |
|---|---|---|---|---|---|
| symp. mexicana | maize | recombination rate (cM/Mb) | 0.385 | 0.341 | 0.428 |
| symp. mexicana | mexicana | recombination rate (cM/Mb) | -0.579 | -0.610 | -0.545 |
| symp. mexicana | parviglumis | recombination rate (cM/Mb) | 0.507 | 0.469 | 0.543 |
| symp. maize | maize | recombination rate (cM/Mb) | -0.066 | -0.117 | -0.014 |
| symp. maize | mexicana | recombination rate (cM/Mb) | 0.011 | -0.038 | 0.061 |
| symp. maize | parviglumis | recombination rate (cM/Mb) | 0.105 | 0.055 | 0.157 |
| symp. mexicana | maize | gene density (coding bp/cM) | -0.262 | -0.309 | -0.212 |
| symp. mexicana | mexicana | gene density (coding bp/cM) | 0.423 | 0.381 | 0.464 |
| symp. mexicana | parviglumis | gene density (coding bp/cM) | -0.385 | -0.428 | -0.341 |
| symp. maize | maize | gene density (coding bp/cM) | 0.050 | -0.003 | 0.101 |
| symp. maize | mexicana | gene density (coding bp/cM) | 0.030 | -0.022 | 0.080 |
| symp. maize | parviglumis | gene density (coding bp/cM) | -0.099 | -0.151 | -0.048 |
