## Supplementary material for "Selective sorting of ancestral introgression in maize and teosinte along an elevational cline": S6 Table

**S6 Table. Diversity within maize and *mexicana* alleles at *inv9f.*** Pairwise diversity ($\pi$) and Watterson's theta ($\theta_W$) for samples clustering by PCA with the maize-allele at the inversion (PC1 $> 0.2$) or *mexicana*-allele at the inversion (PC1 $< 0.4$). Individuals heterozygous for the inversion (or ambiguous in PCA clustering) were excluded. Diversity estimates within the putative inversion region were calculated for each group using the same ANGSD/realSFS pipeline as genomewide diversity estimates. Only subspecies with $> 5$ samples in a cluster were analysed.

| sample | $\pi$ | $\theta_W$ |
|---|---|---|
| mexicana within mexicana inversion cluster | 0.024 | 0.083 |
| maize within mexicana inversion cluster | 0.019 | 0.053 |
| parviglumis within maize inversion cluster | 0.013 | 0.016 |
| maize within maize inversion cluster | 0.011 | 0.022 |
