## Supplementary material for "Selective sorting of ancestral introgression in maize and teosinte along an elevational cline": S7 Table

**S7 Table. Domestication genes and overlap with introgression deserts.**

| gene | phenotype | refs | v4 coordinates | min teosinte introgression into maize | min maize introgression into mexicana |
|---|---|---|---|---|---|
| zagl1 | ear size | Wills et al. 2018 | 1:4959131-5014850 | 0.031* | 0.252 |
| gt1 | prolificacy | Wills et al. 2013 | 1:23605801-23647370 | 0.018* | 0.106 |
| ZmSh1-1 | seed shattering | Lin et al. 2012 | 1:228660490-228705551 | 0.344 | 0.032* |
| tb1 | branching | Doebley, Stec & Gustus 1995, Doebley, Stec & Hubbard 1997, Dong et al. 2019 | 1:270533676-270574776 | 0.023* | 0.041* |
| zfl2 | cob rank | Doebley & Stec 1991, Doebley & Stec 1993, Bomblies & Doebley 2006 | 2:12894091-12937068 | 0.213 | 0.213 |
| pbf1 | storage protein synthesis | Wang, Ueda & Messing 1998 | 2:158122366-158176919 | 0.098 | 0.081 |
| ra2 | inflorescence architecture | Vollbrecht et al. 2005 | 3:12138280-12179065 | 0.132 | 0.053 |
| ba1 | plant architecture | Gallavotti et al. 2004 | 3:185994629-186035264 | 0.257 | 0.191 |
| su1 | starch biosynthesis | Whitt et al. 2002 | 4:43090569-43139167 | 0.542 | 0.006* |
| tga1 | 'naked' grains | Dorweiler et al. 1993 Wang et al. 2005 | 4:46330597-46375118 | 0.062* | 0.028* |
| bt2 | starch biosynthesis | Whitt et al. 2002 | 4:61295575-61341350 | 0.082 | 0.008* |
| ZmSh1-5.1+ ZmSh1-5.2 | seed shattering | Lin et al. 2012 | 5:16630307-16676707 | 0.073 | 0.166 |
| sweet4c | sugar transport and seed size | Sosso et al. 2015 | 5:130767030-130809864 | 0.038* | 0.132 |
| ae1 | starch biosynthesis | Whitt et al. 2002 | 5:172392995-172450415 | 0.291 | 0.06 |
| ra1 | inflorescence architecture | Vollbrecht et al. 2005, Sigmon & Vollbrecht 2010 | 7:113552410-113592937 | 0.16 | 0.148 |

* lowest 5% introgression genomewide ('introgression desert')
